# Functional Imaging for Regenerative Medicine and Tissue-Engineered Constructs: Tracking Vascularization of Tissue-Engineered Small Intestine Scaffolds

**DOI:** 10.1101/2025.09.25.677636

**Authors:** Jared A. Weis, Mani Ratnam Kothamasu, Nildris Cruz-Diaz, Marshall Z. Schwartz, Anthony Atala, Mitchell R. Ladd

## Abstract

Traditional approaches to tissue engineered small intestine (TESI) require implanting a construct in the omentum of the abdominal cavity to allow for vascularization of the construct and tissue maturation. A second operation is then needed to implant the construct in continuity with the intestine. The timing of the second operation has historically been chosen based on pre-determined biological endpoints. Currently, no well-validated method has been used to non-invasively monitor for vascular maturity which would better inform when the second operation should be performed. We hypothesized that photoacoustic imaging (PAI) could serve as a tool to non-invasively monitor regional and temporal changes in vascular maturity in scaffolds implanted in the omentum. For this pilot study, tubular scaffolds used for TESI were fabricated with electrospinning and implanted in the omentum of Sprague-Dawley rats for 1 and 2 months. Scaffolds were imaged with PAI and tissue oxygenation and hemoglobin concentration were quantified. PAI was then correlated with gross observations of vascularization at each time point. PAI was able to determine regional and temporal changes in tissue oxygenation and hemoglobin concentration. Specifically, the oxygenation and hemoglobin concentration of the top wall of the construct showed better vascular maturity compared to the bottom wall. In addition, vascular maturity seemed to improve in the top wall from 1 to 2 months. The bottom wall was not well covered by the omentum and thus did not vascularize as well as the top wall. The PAI findings were confirmed on gross examination of the scaffolds and upon quantification of the histological analysis of endothelial cell density. Thus, PAI may serve as a critical tool for monitoring vascular maturity in TE, specifically within the abdomen. This will be a critical tool in the pre-clinical development and clinical translation of TESI.

## Introduction

Intestinal failure (IF) is the inability to absorb adequate nutrients or fluids to grow and sustain life.^1,2^ The prevalence of IF in the US, including children and adults, is estimated to be 79 patients per million inhabitants.^3^ IF is managed by intravenous calories given through a central venous catheter (parenteral nutrition [PN]), enteral feeds through feeding tubes, and multidisciplinary teams that can oversee the treatment of these complex patients with the hope of promoting intestinal adaptation such that the native intestine will eventually be able to absorb all nutrition through the gut (i.e. enteral autonomy).^4-6^ Treatment is expensive with costs in the US in 2014 estimated to be $2.3 billion.^4,7,8^ More importantly, these therapies have numerous morbidities such as central venous catheter complications, sepsis, diarrhea, intestinal-failure-associated liver disease, and death.^4,5,9,10^ The all-cause mortality of IF ranges from 7-16% and is primarily driven by central venous access complications and liver failure from chronic PN.^5,11-14^ Tissue-engineered small intestine (TESI) has emerged as a potential treatment strategy for IF.

Previous efforts to create a TESI have focused on the epithelium and nutrient absorption with less attention on the critical requirement of peristalsis which is vital for nutrient absorption.^15-29^ We have focused our initial work on the intestinal muscular layer and have successfully created a tubular scaffold with aligned fibers that induces human intestinal smooth muscle cell (hiSMC) alignment *in vitro* which is a good candidate for developing a tubular tissue construct with organized muscle that may be capable of peristalsis once implanted *in vivo*.^2^ In the rodent animal model to test TESI for tissue formation and function, the seeded construct is first implanted in the omentum to allow the muscle cells to form tissue and to obtain a blood supply. Four to six weeks after initial implantation, the constructs are placed in continuity with the intestine. This fixed interval time point is chosen based on convenience and general assumptions of the time course of inflammation without any direct knowledge of how each individual construct has matured prior to the second operation. The inability to functionally query the changes of an implanted construct over time thus represents a significant gap in the TESI field and is limiting to the advancement of tissue-engineering and regenerative medicine in general.

New methods of *in vivo* noninvasive imaging-driven assessment of implanted constructs are therefore required, capable of reporting on construct vascularity – a critically important criterion to construct success. Photoacoustic imaging (PAI) has emerged as a method that has significant untapped potential to fill this need. PAI generates image contrast through ultrasound detection of optical absorption events. By incorporating spectrally selective near-infrared imaging, oxy- and deoxyhemoglobin can be separately visualized and quantified using computational spectral unmixing methods to allow for spatial assessment of tissue blood flow and oxygen saturation. While PAI has been used to monitor scaffolds in tissue engineering (TE) previously, its use has been limited.^30-34^ Moreover, PAI has never been used to track vascular maturation of scaffolds within the abdominal cavity, a unique requirement for TESI. We hypothesized that PAI could be used to measure functional maturation in TESI construct vascularization *in vivo* over time thereby informing in future work when the constructs are ready to be placed in continuity with the small intestine. In this pilot study, we demonstrate the use of noninvasive functional vascular imaging of tissue oxygenation and hemoglobin content to track temporal changes in construct vascular maturity within the abdominal cavity (**Figure 1**).

**Figure 1.**
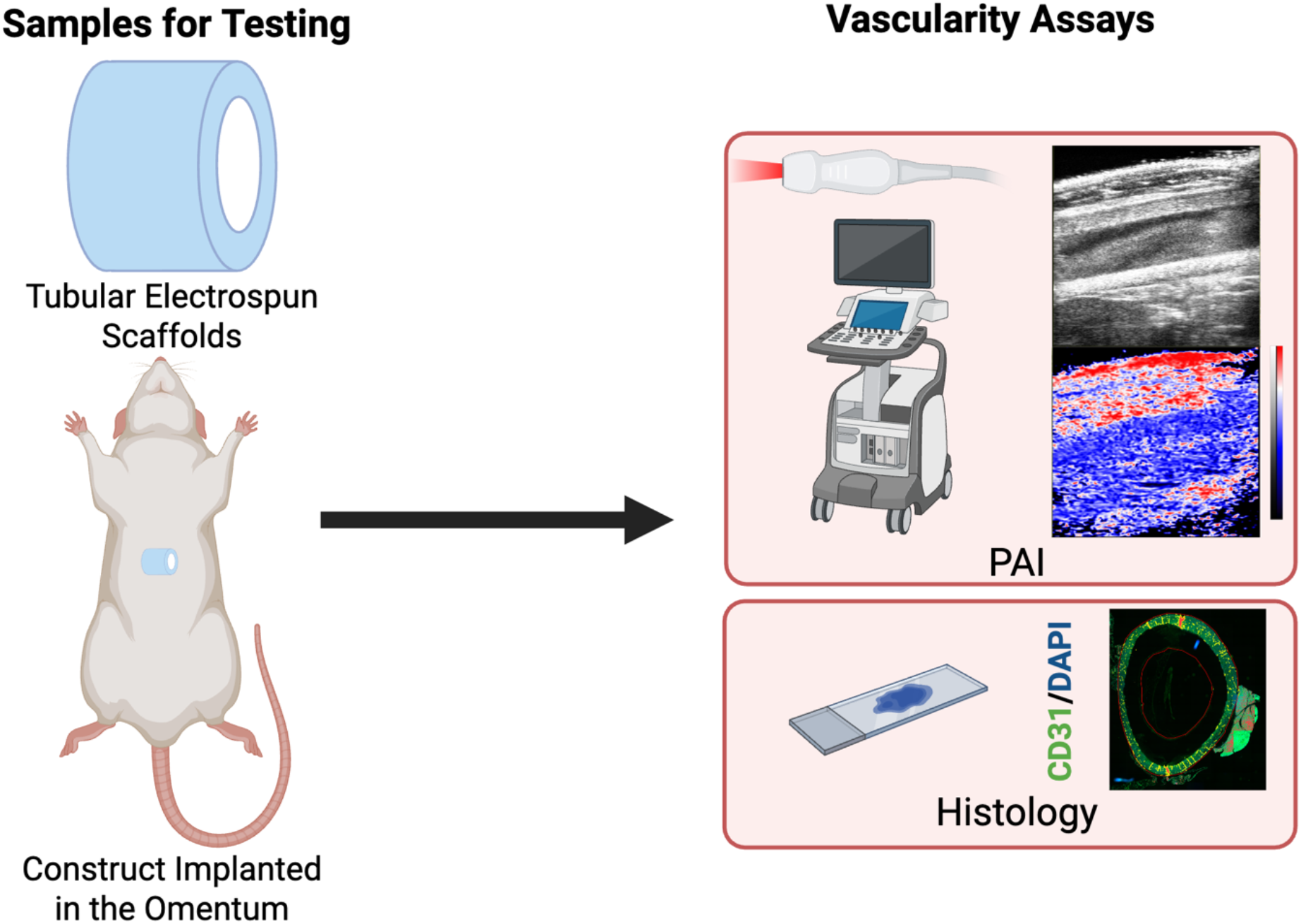
Study design and overview. Tubular scaffolds are fabricated with electrospinning using a combination of polycaprolactone and type I collagen. The constructs are then implanted into the omentum of Sprague-Dawley rats. At 1- and 2-months the vascularity of the constructs is evaluated using photoacoustic imaging and correlated to end-point histological staining for CD31 a vascular endothelial marker.

## Methods

### Materials List

Pluronic F-127 (Sigma Aldrich, cat# P24434), Polycaprolactone (PCL, Evonik/Lactel absorbables, cat#B6003-2 and Sigma 24980-41-4, cat# 24980-41-4, inherent viscosity range 1.0-1.9 dL/g cat#900820), calf skin type I collagen (Elastin Products Company, cat#c857), hexafluoroisopropanol (HFIP, Sigma Aldrich, cat#105228), 1-ethyl-3-(3-dimethylaminopropyl)carbodiimide hydrochloride (EDC, Thermofisher Scientific, cat#22981), N-Hydroxysuccinimide (NHS, Sigma Aldrich, 130672).

### Scaffold Fabrication

Scaffolds were fabricated as previously described in Ladd *et al*.^2^ In brief, tubular scaffolds consisting of a one-to-one blend of PCL and type I collagen (with inner diameter of 6 mm were fabricated using electrospinning. The polymer/collagen solution was 18% wt/vol and dissolved in HFIP. Prior to electrospinning the polymer solution, 3 ml of a 20% wt/vol Pluronic F-127 solution in deionized water was used to coat the mandrel so that the final scaffold would be easier to remove from the mandrel. Then, the PCL/collagen solution was spun at a flow rate of 10 ml/hr through a 16-gauge blunt tip needle, with a needle to mandrel distance of 15 cm and needle tip voltage of 20 kilovolts. The mandrel rotation speed was 11,000 RPMs to promote alignment of electrospun fibers in the circumferential direction. The scaffolds were allowed to dry and then crosslinked with EDC and NHS for 24 hours. After crosslinking, the scaffolds were rinsed in deionized water, submerged in 100% ethanol to wet the Pluronic and remove the scaffold from the mandrel, frozen at -80^º^C for 2 hours, and then lyophilized. After lyophilization, scaffolds were stored at -80^º^C until use. These scaffolds were previously characterized in Ladd *et al*.^2^

### Scaffold Omental Implantation

All animal surgeries and imaging studies were approved by the Institutional Animal Care and Use Committee (Protocol A24-063). Sprague-Dawley rats. between the ages of 8 weeks and 6 days and 9 weeks and 1 day old at the time of scaffold implantation were used for all studies.

Prior to implantation, the scaffolds were cut to 4 cm in length. The day prior to surgery, the scaffolds were sterilized by placing them in 70% ethanol for 2 hours followed by three 20-minute rinses in sterile PBS under a laminar flow hood. The constructs were then left under UV light for at least 2 hours after which they were placed in the 37^º^C incubator overnight to acclimate to body temperature prior to implantation. After sterilization, animals were appropriately anesthetized after which a midline laparotomy was performed. The scaffold was then wrapped in the omentum and the omentum was secured in place on the right aspect of the construct using a simple 4-0 vicryl suture. The abdomen was then closed in layers in the usual fashion. Animals were then survived until their time point at 1- or 2-months post-surgery. At each time point, the rats were anesthetized for photoacoustic imaging and then were euthanized. After euthanasia, gross images were taken of the constructs to compare to imaging and then the constructs were explanted for histological analysis.

### Photoacoustic Imaging

Rats were anesthetized using isoflurane delivered in 100% oxygen (100% FiO2) and high frequency anatomical US images were acquired with co-registered PAI images acquired at 750 nm and 850 nm spectral excitation wavelengths using a Vevo F2 LAZR-X (FUJIFILM VisualSonics, Inc., Toronto, ON, Canada). This enabled high-resolution measurements of oxyhemoglobin and deoxyhemoglobin concentration throughout the abdomen of the rats. 2D images were centered at the construct midplane in the transverse orientation. 3D images were acquired based on 2D slices acquired at multiple locations perpendicular to the imaging plane and used to build a 3D volume with US imaging transducer location controlled automatically via a motorized controller. Construct oxygenation was assessed based on the calculation of oxygen saturation using PAI measurements of oxyhemoglobin and deoxyhemoglobin concentrations.Oxygen saturation (sO2) was calculated as the ratio of oxyhemoglobin to total hemoglobin (HbT), performed using vendor-supplied analysis tools. Regions-of-interest (ROIs) were drawn to indicate both the top and bottom walls of the construct within the abdomen and used to measure average construct blood oxygen saturation and total hemoglobin in the top half of the scaffold compared to the bottom half for each rat. Mean sO2 and HbT measurements were recorded for ROIs drawn from the anatomical US images to designate the top and bottom walls of the construct in 2D analysis of the construct midplane. Following initial imaging assessment under 100% FiO2, a breathing gas challenge was used in which the carrier inhaled gas was changed to medical air (21% FiO2) and imaging measures were repeated.

### Histological Analysis

At the time of euthanasia, gross images were obtained, and the constructs were removed from the omentum. A center section of the construct was excised for histology and placed in 10% neutral-buffered formalin and fixed for 2-3 days. Then it was transferred to 70% ethanol until it was further processed and paraffin embedded. The constructs were embedded to obtain both cross-sections. Sections were cut at 5 microns and then stained with hematoxylin and eosin (H&E). These slides were imaged on the E1000 Dx Digital Pathology Solution at 200X with the entire tissue sections being captured allowing for digital changes in magnification (Epredia).

Sections also underwent immunostaining to evaluate for vascular endothelial cells using an antibody for CD31 (rabbit anti-CD31 [EPR17259, Abcam, ab182981, 1:1000 dilution). The slides underwent antigen retrieval with EDTA at pH 9 prior to staining. The secondary antibody used was donkey anti-Rabbit IgG 488 (Biotium, 20015, 1:1000 dilution) and the slides were counterstained with DAPI. Afterwards, the slides were imaged on the Evident Slideview VS200 (Evident) at 200X. The entire tissue section was imaged at 200X which then allowed for digital changes in magnification. The images were then analyzed using QuPath.^35^ Specifically, the images were opened, and the tissue area was manually outlined to designate a region of interest. Positive fluorescent area was designated above a threshold of 2500 within the tissue region of interest. The threshold was chosen as the value at which QuPath was selecting the true positive staining. The percent positive area was then calculated as the positive threshold area divided by the total tissue area. This percentage was then taken as the approximate value of positive CD31 staining with higher percent area positivity denoting higher microvascular density.

### Statistical Analysis

All data are shown as a mean with standard deviation. All data were analyzed with unpaired or paired two-tailed t-tests (as specified) or one-way ANOVA with post-hoc Tukey’s test as appropriate with significance taken as a p value less than 0.05.

Data were analyzed using GraphPad Prism software.

## Results

### Scaffold Implantation

Surgeries for a total of 12 animals, 6 animals each in the 1-month and 2-month time points, were without issues. Three total animals had to be euthanized due to humane endpoints and at necropsy were found to have bowel obstructions where the bowel slipped into the lumen of the scaffold. This was thought to have occurred because of the 4 cm length of scaffold used and the relatively young (approximately 9-weeks old) age and low weight (approximately 250-300 g) of the rats used, with a less developed omentum that was not quite large enough to fully wrap the entire scaffold. Thus, there were 4 rats for the 1-month time point and 5 rats for the 2-month time point. All remaining rats survived until their planned time point.

### Photoacoustic Imaging

At the 1- and 2-month time points, PAI demonstrated that the regions of the implanted scaffold were well-oxygenated (**Figure 2**). Of interest, the top wall of the implanted scaffolds at both 1- and 2-months were found to have more oxygen saturation compared to the bottom wall of the scaffold. When quantifying the oxygen saturation from PAI at 100% inhaled oxygen concentration between 1- and 2-months, there was a significantly higher oxygen concentration in the top wall at both time points compared to the bottom wall (**Figure 3**). There was no increase in tissue oxygen saturation in the top wall of the scaffolds from 1- to 2-months. Similarly, there was no increase in tissue oxygen saturation in the bottom wall of the scaffolds from 1- to 2-months.

**Figure 2.**
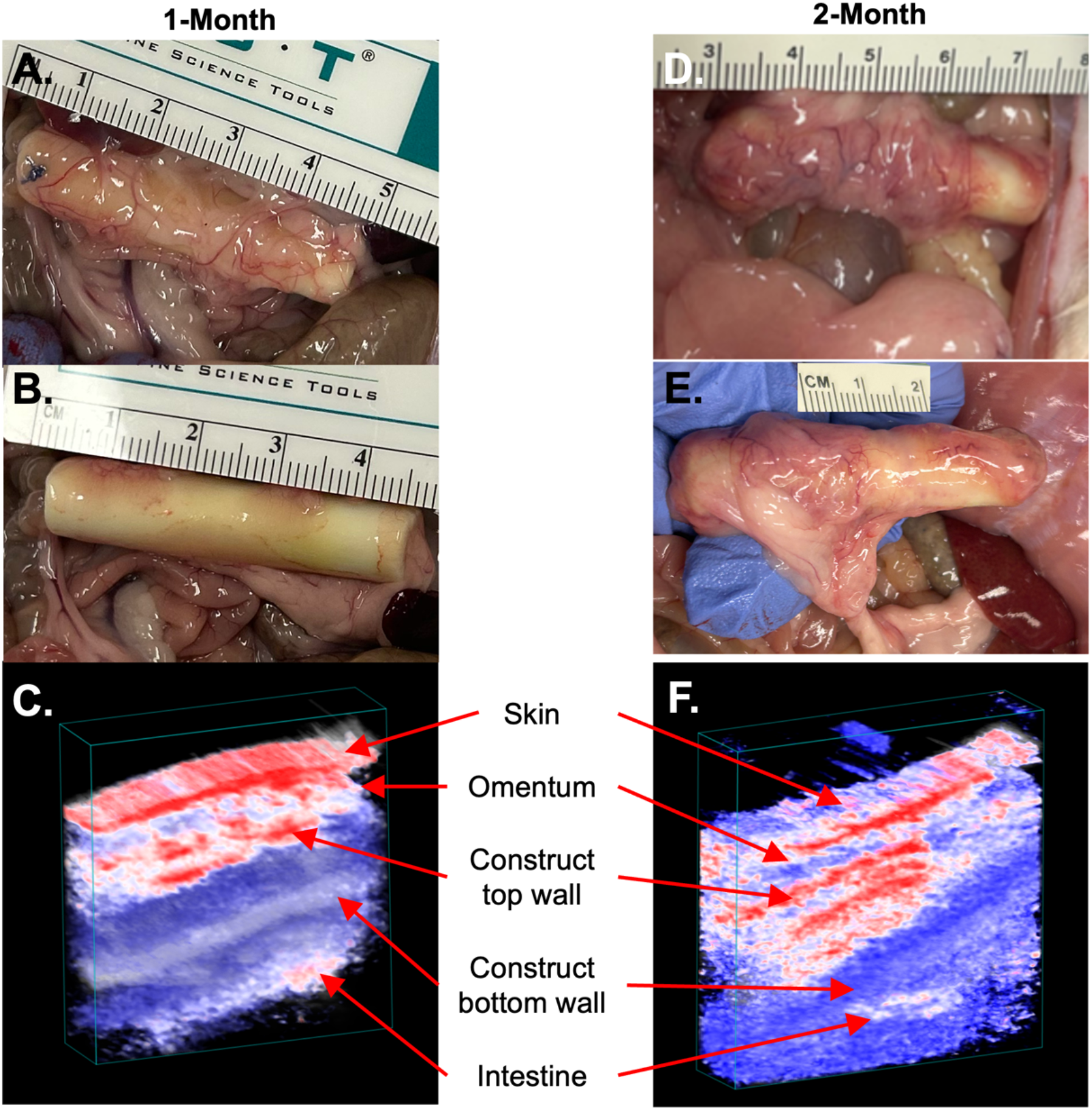
Gross evaluation of implanted scaffolds and correlation to photoacoustic imaging reconstructions at 1- and 2-month time points. **A.** *In situ* depiction of a representative implanted scaffold on the anterior/top surface at 1 month. **B**. *In situ* depiction of the same implanted scaffold on the posterior/bottom surface at 1 month. Note that the posterior side was not as well covered by omentum and thus not as well vascularized. **C**. 3D reconstruction of the photacoustic imaging co-registered with ultrasound at 1 month. Note that the top wall of the construct has more vascularity than the bottom wall as noted grossly at necropsy. **D**. *In situ* depiction of a representative scaffold on the anterior surface at 2 months. **E**. *In situ* depiction of the same scaffold on the posterior surface. While somewhat better covered, note that the posterior aspect of the scaffold is not as well vascularized. **F**. 3D reconstruction of the photoacoustic imaging co-registered with ultrasound at 2 months. Again, note that the vascularization is less on the posterior wall consistent with what was seen grossly at necropsy.

**Figure 3.**
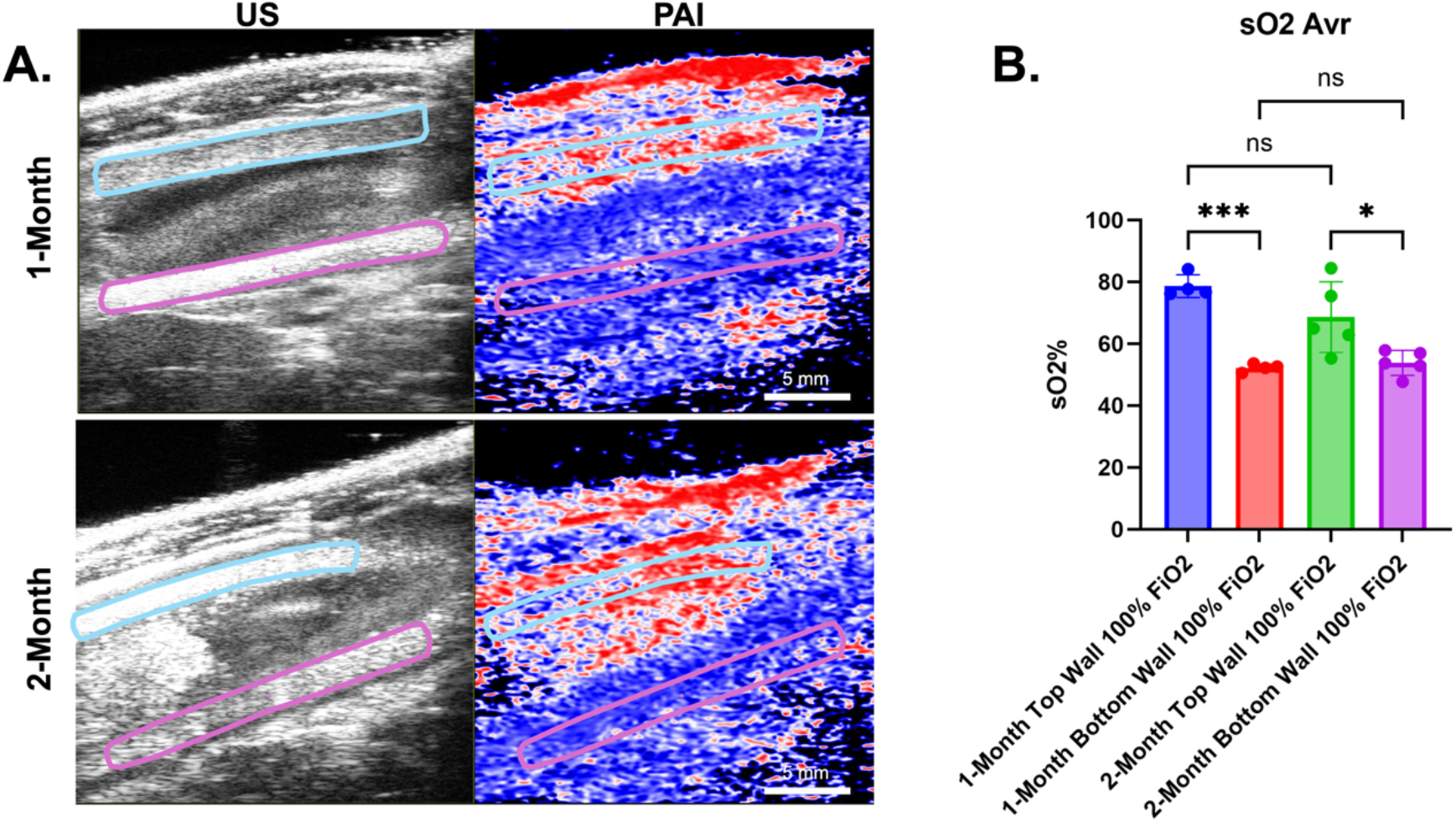
**A.** Representative ultrasound and photoacoustic images of implanted scaffolds at 1- and 2-month time points when the animal was inhaling 100% oxygen. Like Figure 2, note the oxygenation in the top wall of the scaffold compared to the bottom wall. The blue box indicates the top wall region of interest, and the purple box indicates the bottom wall region of interest on the ultrasound images. **B**. Oxygen saturation quantification of the top and bottom walls of the construct at 1 month compared to 2 months. There were statistically significant differences in tissue oxygenation between top and bottom walls at both time points. There were no significant differences in tissue oxygen saturation in the top wall from 1 to 2 months nor the bottom wall from 1 to 2 months.

Next, the concentration of inhaled oxygen was changed from 21% FiO2 to 100% FiO2 in a breathing gas challenge to further validate PAI detection of oxygen saturation changes within the construct responsive to systemic oxygen saturation changes (**Figure 4**). In doing this, there was a clear visual difference overall in the amount of oxygen saturation within the construct at 100% FiO2 as compared to 21% FiO2 (**Figure 4A**). This was specifically apparent within the top wall at 1- and 2-months, which showed a statistically significant decrease in oxygen saturation in the top wall of the scaffold when decreasing in inhaled oxygen concentration from 100% to 21% (**Figure 4B** and **C**). Because the bottom wall was less vascularized, a similar reduction in oxygen saturation was not seen in the bottom wall of the construct when decreasing from 100% to 21% oxygen. This confirmed the ability of PAI to detect functional vasculature within the scaffold when present, with changes in inhaled oxygen concentration leading to changes in the oxygen concentration in the scaffold on PAI.

**Figure 4.**
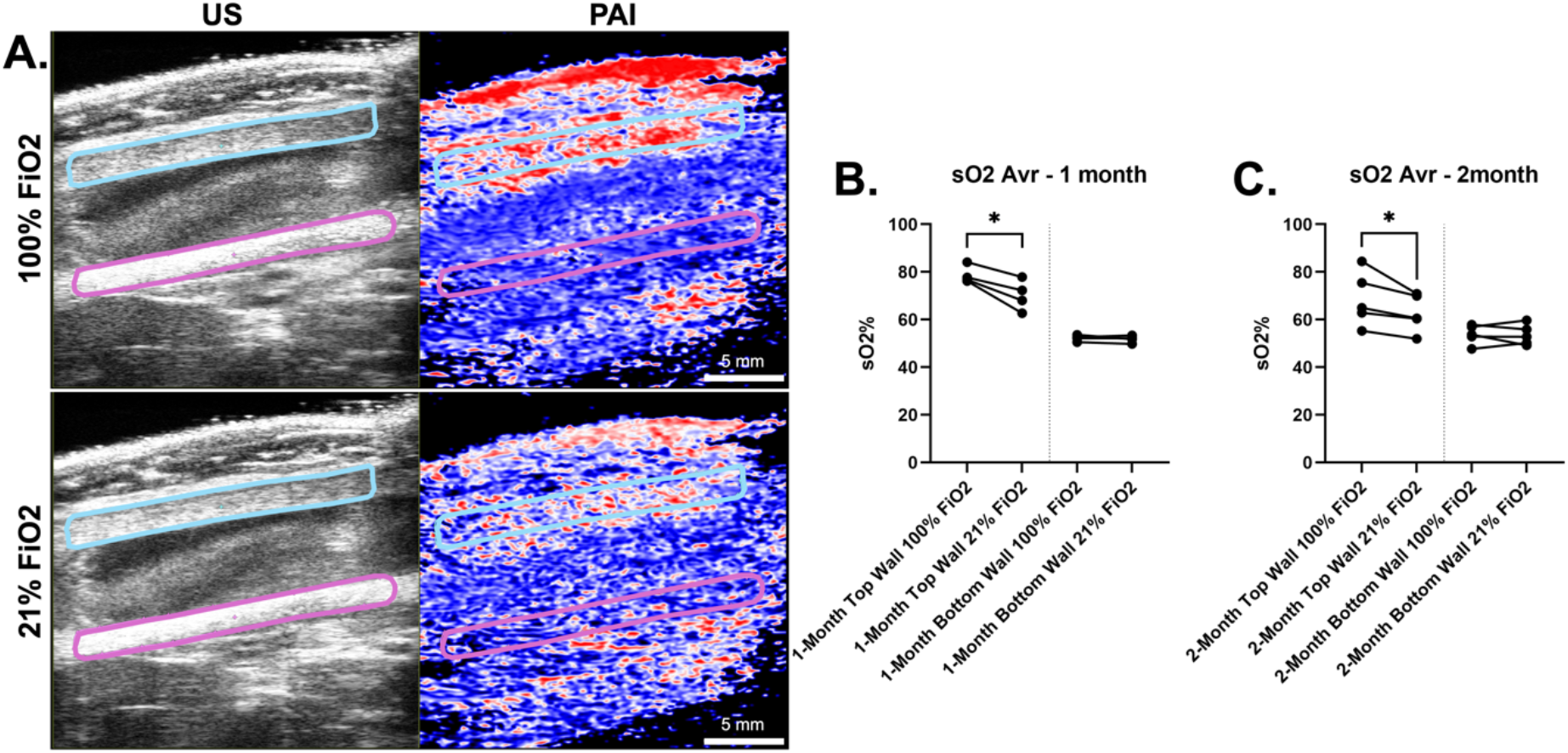
**A.** Representative photoacoustic images of the oxygenation of the implanted construct at 1 month when the animal is exposed to 100% oxygen vs. 21% oxygen concentration. When comparing 100% to 21% there are more areas in red representing higher oxygenation, especially along the top wall of the scaffold compared to the bottom. The blue box indicates the top wall region of interest, and the purple box indicates the bottom wall region of interest on the ultrasound images. **B**. Quantification of oxygen saturation at 1 month between 100% and 21% inhaled oxygen between the top walls and bottom walls of the scaffold. A two-tailed paired t-test was used. **C**. Quantification of oxygen saturation at 2 months between 100% and 21% inhaled oxygen between the top walls and bottom walls of the scaffold. A two-tailed paired t-test was used.

Finally, we examined total hemoglobin content in the implanted scaffolds at both 100% and 21% inhaled oxygen using PAI (**Figure 5**). Given the relatively low amount of oxygen saturation and thus vasculature in the bottom wall of the scaffold, we evaluated only the top wall of the scaffold. There was an increasing trend in total hemoglobin content in the top wall from 1-month to 2-months both at 21% and 100% inhaled oxygen, however, only the increase at 21% FiO2 was statistically significant (**Figure 5B**). This was consistent with our gross observations which revealed increased vascularization of the 2-month scaffolds in general compared to the 1-month scaffolds. Again, this demonstrated the ability of PAI to detect regional and temporal changes in not only tissue oxygen saturation, but also total hemoglobin content which is a surrogate measure for vascularization.

**Figure 5.**
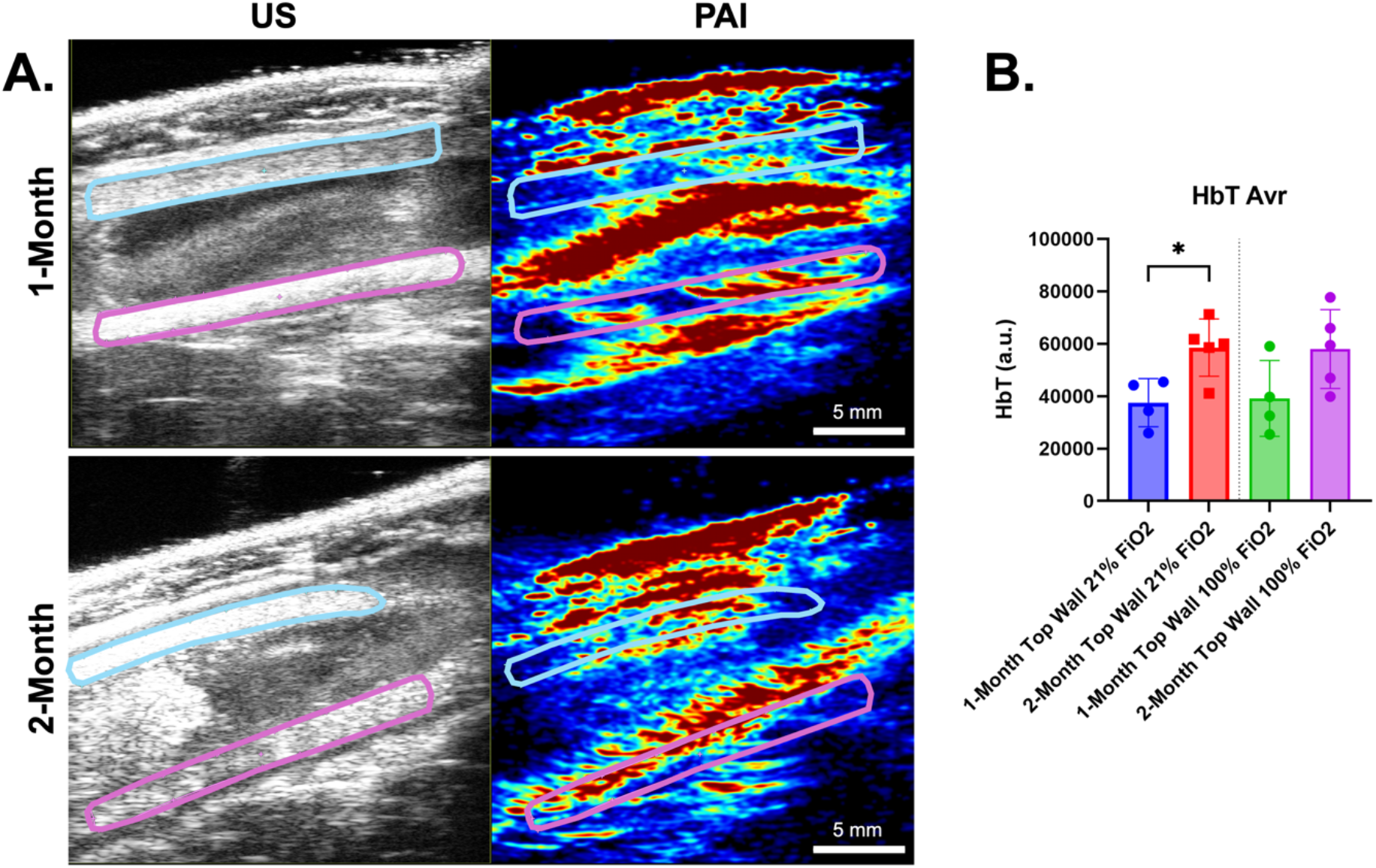
**A.** Representative photoacoustic images demonstrating total hemoglobin content at 1 and 2 months when the rats are inhaling 21% oxygen. The blue box indicates the top wall region of interest, and the purple box indicates the bottom wall region of interest on the ultrasound images. **B**. Quantification of total hemoglobin content of the top wall at 1 and 2 months at either 100% or 21% inhaled oxygen concentration. Note that in both cases the 2-month time point has a higher total hemoglobin concentration, however the difference only reached statistical significance during the 21% concentration of inhaled oxygen. A two-tailed unpaired t-test was performed.

### Gross Examination

Gross observations at necropsy revealed that the omentum was covering the top or anterior surface of the scaffolds better than the bottom or posterior side of the scaffold. This was consistent with the PAI findings. Because of this, each animal served as an internal control demonstrating that we could differentiate the vascularization seen on the top where it was better compared to the bottom. When carefully examining the gross surfaces, there seemed to be more vascular ingrowth and angiogenesis at 2 months compared to 1 month on average. In addition to the vascularization of the scaffold, we could easily identify the other layers of the abdomen including the skin and omentum, and the underlying intestines (**Figure 2**).

### Histology

Staining and analysis of the scaffolds after explantation from the omentum demonstrated surrounding omental and inflammatory tissue. In addition, there were areas of vascular ingrowth noted on hematoxylin and eosin (**Figure 6**). The CD31 staining showed that endothelial cells were contained within the surrounding tissue as well as within the scaffold (**Figure 6B, C, E**, and **F**). Of note, the amount of CD31 positive staining was found to be more prevalent and deeper into the scaffold in the 2- month samples as compared to the 1-month samples. When the samples were quantified, there was an increase in positive staining area of CD31 at 2-months compared to 1-month.

**Figure 6.**
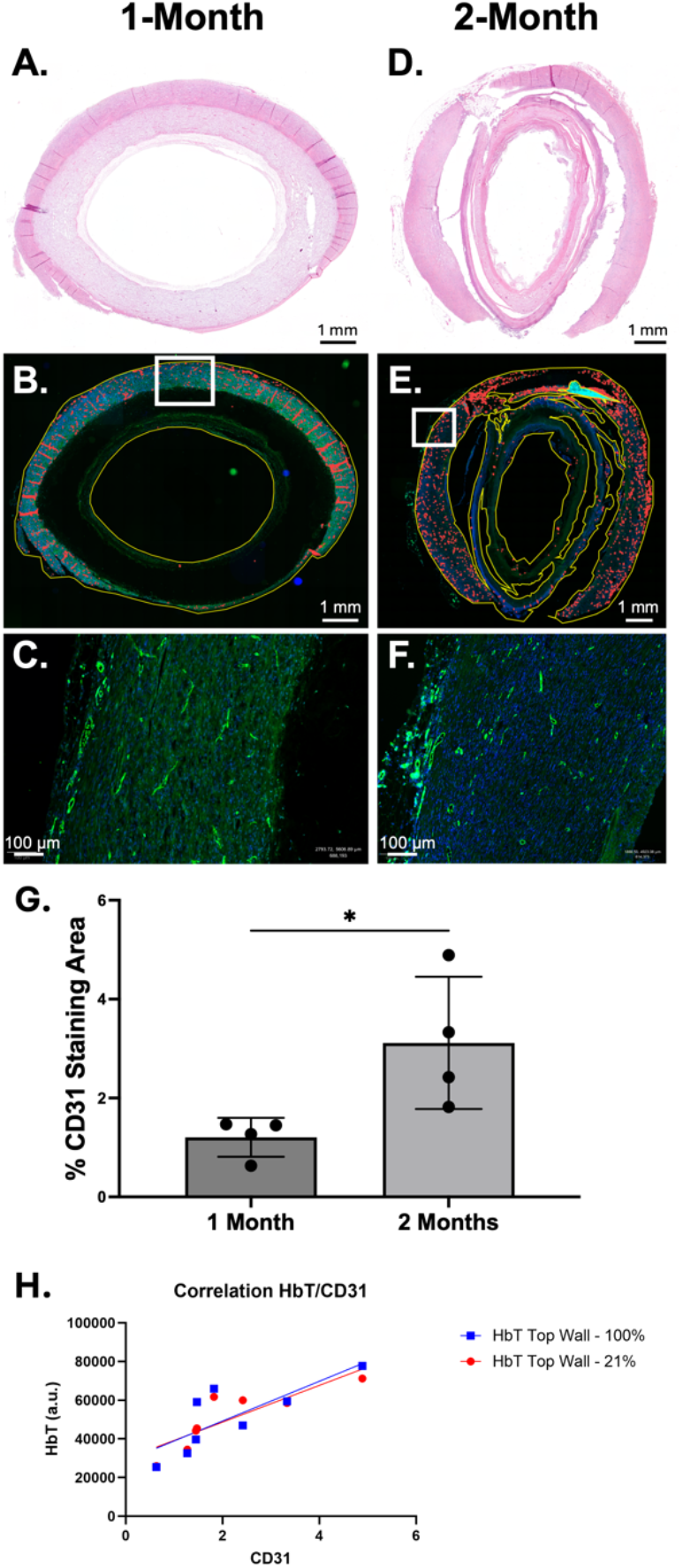
**A.** A representative hematoxylin and eosin stain of a 1-month time point scaffold. The tissue surrounds the scaffold and is starting to grow into it. **B**. CD31 immunostaining of the same representative sample along with the segmentation used to determine total tissue area and positive staining area. **C**. A magnified image of the CD31 staining showing vascular endothelial cells within the wall of the scaffold. **D**. A representative hematoxylin and eosin stain of a 2-month time point scaffold. **E**. CD31 immunostaining of the same representative sample along with the segmentation used to determine total tissue area and positive staining area. **F**. A magnified image of the CD31 staining showing vascular endothelial cells within the wall of the scaffold. **G**. Quantification of percent positive area staining with CD31 at 1 and 2 months. There was a statistically significant increase in CD31 positive staining area at 2 months consistent with the overall increase in tissue oxygenation and total hemoglobin noted. **H**. Pearson’s correlation showed a statistically significant correlation between total hemoglobin in the top wall and CD31 staining density at 100% (Pearson r=0.7911, p=0.0194) and 21% (Pearson r= 0.8499, p=0.0075) FiO2.

## Discussion

This pilot study demonstrates the feasibility of PAI to monitor TE scaffold or construct vascular maturity over time. We show that PAI can detect changes in both tissue oxygenation and hemoglobin content. We internally validated our findings through the use of a breathing gas challenge wherein we changed the inhaled oxygen concentration from 100% to 21% and detected a clear decrease in tissue oxygenation, confirming that PAI detects functional vasculature within the wall of the scaffold. Moreover, the differences detected by PAI correlated well with our observations both grossly and histologically. The top wall of the implanted constructs was found to have more oxygenation and hemoglobin than the bottom wall on PAI and this difference in vascular ingrowth was evident on necropsy of the animals. In addition, PAI measured total hemoglobin concentration increased from 1-month to 2-months in the top wall of the scaffold suggesting an increase in vascularity. This was confirmed histologically with staining of vascular endothelial cells with CD31 and quantification of microvessel density. To our knowledge, this is the first report of using PAI to monitor changes in vascularization of TE scaffolds in the abdomen over time. PAI is feasible and reliable in detecting tissue oxygenation and hemoglobin content in implanted constructs within the abdomen and can distinguish the scaffold wall from surrounding tissues such as the omentum and intestine. Finally, histological quantification of microvessel density correlates well with PAI imaging, validating imaging findings.

While PAI was able to detect functional vasculature within the wall of the scaffold, the lack of increase in tissue oxygenation within the top wall from 1 to 2 months was initially surprising as we observed on gross inspection that there appeared to be more vascular ingrowth at 2 months. For the bottom wall, the lack of increase in vascularity was consistent with our gross observations of the scaffolds at both time points. However, upon analysis of total hemoglobin, we deduced that vascularity was increasing and that tissue saturation was generally the same as saturation is not dependent on total hemoglobin but is the percent of oxygenated hemoglobin. Regardless, the results demonstrated that PAI can detect regional differences and changes in scaffold tissue oxygenation over time. The breathing gas challenge indicated that the scaffolds had developed blood flow, presumably through angiogenesis, and that these vessels were responsive to changes in inhaled oxygen concentration. Histological analysis confirmed that there was an increase in microvessel density from 1- to 2-month time points validating the PAI findings of total hemoglobin increase from 1 to 2 months.

While the development of PAI began in the 1960s, the modern use of PAI for *in vivo* imaging did not begin until the 1990s.^36^ The first use of PAI in tissue engineering was reported in 2009 by Nam *et al*. who evaluated the vascularization of hydrogels that were implanted subcutaneously in rats. Prior to imaging, however, Nam *et al*. removed the constructs and performed PAI *ex vivo*. Since then, others have used PAI to evaluate implanted scaffolds or constructs, but these have been implanted subcutaneously or in the abdominal wall and not within the abdominal cavity.^31-33^ Despite this early work, the use of PAI has not accelerated in TE and seems underutilized.

The motivation to use PAI in TESI was born out of experience with the *in vivo* model. Specifically, TESI constructs are placed into the omentum to mature and develop a blood supply. Then, after a pre-determined time point based on the known time course of inflammation and angiogenesis, the construct is then placed into continuity with the intestine during a second surgery using the omental flap as the blood supply for the construct. However, in our experience, there are times upon performing the second operation that the construct is not yet well-vascularized or is incompletely wrapped by the omentum. This problem spurred the motivation for this work. Non-invasive monitoring of the implanted constructs would allow us to monitor construct vascular maturity to avoid this situation. Alternatively, in the case of a construct in which the omentum had separated from the construct, combining the use of US and PAI would allow early identification of this problem and alert to the need to perform a reoperation to better wrap the omentum around the construct.

The study did have several limitations. First, the overall sample size was small as this was a proof-of-concept pilot study. Second, the sample size was further decreased due to early mortality in the animal model. As discussed, this mortality was likely due to the use of younger rats with less robust omentum in combination with scaffolds that were too long. In future work we will use older rats with better developed and larger omentum along with shorter length scaffolds for implantation. Third, we did not orient scaffolds with ink or other landmark fiducials when they were removed from the omentum. This limited direct histological correlation to imaging results as we are unable to conclusively comment on top versus bottom wall from a histology perspective. Fourth, we performed staining of single sections to perform histological validation assessment, however this could be improved by performing serial sectioning with subsequent quantification. Lastly, we examined only two relatively short time points in this pilot study, but future work will examine more and longer construct maturation time points.

While this pilot study shows significant promise, we used a scaffold with a single material composition. Future work will investigate scaffolds composed of short- and long-term biodegrading materials to determine differences in vascularization between material types and further validate the imaging assessment method. In addition, as we progress in developing TESI, we will begin to include cell-seeded constructs and will monitor them for vascular maturity.

## Conclusions

PAI is a promising imaging modality for monitoring *in vivo* vascular maturity of tissue engineered scaffolds and constructs. We have demonstrated the feasibility of PAI to monitor regional and temporal changes in vascular maturity of tubular scaffolds implanted in the omentum of the abdominal cavity. PAI data were then validated with gross observations and histological quantitative analysis of vascularization. Future work will expand upon this pilot study to evaluate larger sample sizes with longer time points and diverse biodegradable materials with varying degradation rates. PAI holds promise as a critical tool in pre-clinical TE development and ultimate translation of TE technologies to patients.

## Acknowledgements

The authors would like to thank the Wake Forest Institute for Regenerative Medicine for its ongoing support.

We would like to acknowledge the Preclinical Ultrasound and Photoacoustic Imaging Core of Wake Forest School of Medicine supported in part by the Wake Forest Clinical and Translational Science Institute (NIH NCATS UL1TR001420), and Hypertension and Vascular Research.

## Authorship Contributions

JW: Conceptualization, data curation, formal analysis, investigation, methodology, project administration, resources, software, validation, visualization, writing original draft, writing review and editing.

MRK: Data curation.

NCD: Data curation.

MZS: Conceptualization, resources, writing review and editing.

AA: Conceptualization, funding acquisition, writing review and editing. MRL: Conceptualization, funding acquisition, data curation, formal analysis, investigation, methodology, project administration, resources, supervision, validation, visualization, writing original draft, writing review and editing.

